# Structural variants contribute substantially to complex trait heritability

**DOI:** 10.64898/2026.02.28.708732

**Authors:** Dat Thanh Nguyen, Alexey A. Shadrin, Nadine Parker, Julian Fuhrer, Nam S. Vo, Anders M. Dale, Ole A. Andreassen, Oleksandr Frei

## Abstract

Despite accumulating evidence that structural variants (SVs) exert disproportionate functional effects, their genome-wide contribution to complex trait heritability has not yet been systematically quantified. Here, we introduce MiXeR-SV, a tool that integrates long-read-derived SVs from existing reference catalogs with genome-wide association summary statistics to quantify SV heritability and enrichment. Applying MiXeR-SV to 105 complex traits, we identify 31 traits with significant enrichment (Bonferroni-corrected *P* < 4.8 × 10^−4^), with SVs explaining up to 32% of total heritability, despite comprising only 0.6% of the analyzed variants. Enrichment is most extensive in hematological, metabolic/biomarker, and cancer traits, trait-specific in neuropsychiatric and cardiometabolic phenotypes, and observed across all anthropometric traits, albeit with modest effect sizes, consistent with their highly genetic architecture. These findings are robust across two independent SV reference panels built from complementary long-read and graph-based variant catalogs, with 93.5% of significant traits consistent between technologies. Furthermore, incorporating SV architecture into heritability models consistently increases total heritability estimates in enriched phenotypes, with SV heritability proportions correlating with the gap between SNP-based and twin heritability estimates across traits. Cross-ancestry replication in Biobank Japan confirms enrichment patterns. Our quantification of SVs contributions to genetic architectures has significant implications for genetic prediction and fine-mapping of human complex traits and common diseases.

## 1 Introduction

Genome-wide association studies (GWAS) have identified tens of thousands of common and rare genetic variants associated with complex human traits and diseases, yet common genetic variants explain only a fraction of trait heritability (*h*^2^)[1, 2]. Recent whole-genome sequencing analyses show that rare variants (minor allele frequency, MAF*<*1%) explain approximately 20% of trait heritability[3], but these estimates predominantly capture single nucleotide polymorphisms (SNPs) and small indels while overlooking larger structural variants (SVs). Molecular evidence suggests that SVs may contribute substantially to complex trait heritability, as SVs are highly enriched among lead expression quantitative trait loci (eQTLs), indicating a disproportionate regulatory impact[4, 5]. Yet their contribution to *h*^2^ has not been systematically quantified, representing an important gap in our understanding of complex trait genetics.

SVs are genomic alterations typically defined as ≥50 base pairs, including deletions, duplications, and complex rearrangements such as translocations [6]. They represent a major source of genetic diversity in human populations[6, 7]. Each human genome harbors approximately 20,000 SVs compared with millions of SNPs, yet SVs affect a comparable total number of nucleotides per genome and exhibit disproportionate functional effects[4, 5, 8, 9]. Molecular analyses demonstrate that an SV is 28 to 54 times more likely to modulate gene expression than an SNP or indel[5]. Despite this outsized per-variant impact, SVs were not well captured by short-read sequencing, limiting possibility for their imputation from traditional SNP arrays, leaving their aggregate heritability contribution unmeasured.

Two technical barriers have prevented comprehensive SV heritability estimation. First, long-read sequencing technologies such as Oxford Nanopore Technologies (ONT) provide superior SV detection performance [10, 11]; however, their substantial per-sample cost limits the availability of population-scale cohorts required for robust heritability estimation. Large-scale genomic initiatives have consequently relied on genotyping arrays or short-read sequencing technologies[12–14]. For instance, the UK Biobank (UKBB) whole-genome sequencing effort, which represents one of the largest population genomics resources with 490,640 participants, employed exclusively short-read Illumina technology and consequently captured only a limited fraction of the complete SV landscape[14]. While recent SV imputation panels built from long-read sequencing now enable recovery of common SVs in biobank cohorts[15], they do not address the question of SV heritability even when SV genotypes are available. This is because existing frameworks such as linkage disequilibrium score regression (LDSC)[16, 17] and restricted maximum likelihood (REML) approaches[18] were developed for SNP-based analyses and lack the ability to to partition heritability contributions from structural variations. Although annotation-based extensions of these frameworks have been applied to diverse genomic features[17], none explicitly models the distinct effect size distribution of SVs or integrates comprehensive long-read-derived SV catalogs with summary-level GWAS data, which represents an unresolved challenge in complex trait genetics.diseases

Here, we introduce MiXeR-SV, a summary statistics-based method that quantifies SV contributions to trait heritability by integrating comprehensive long-read SV reference panels with large-scale SNP-based GWAS summary statistics. Leveraging the 1000 Genomes Project (1KGP) resource comprising both whole-genome short-read sequences and 1,019 ONT long-read samples[19, 20], we construct ancestry-specific unified SNP-SV reference panels for European (EUR) and East Asian (EAS) populations. We estimate partitioned heritability within each variant class while modeling MAF and linkage disequilibrium (LD) dependent architecture, and calculate fold-enrichment of heritability explained by SVs relative to genome-wide expectations. By jointly modeling common SNPs and SVs, our method estimates combined variant-based heritability 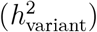. Although 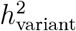 remains a lower bound on narrow-sense additive heritability due to incomplete coverage of rare variants, it provides a broader accounting of additive genetic variance than conventional SNP-based heritability estimates and offers a principled step toward resolving missing heritability.

Applying MiXeR-SV to 105 complex traits from predominantly EUR ancestry cohorts, we identify 31 traits with genome-wide significant SV enrichment, with fold-enrichment ranging from 6.6 to 61.5. We verify our findings using an independent reference panel derived from 3,202 1KGP samples genotyped through the Human Genome Structural Variation Consortium phase 3 (HGSVC)[21] as a technical validation. Cross-ancestry replication in Biobank Japan (BBJ)[12] further confirms significant SV enrichments. Our framework enables scalable SV heritability quantification from existing GWAS summary statistics without requiring population-scale long-read sequencing, with implications for polygenic risk prediction, fine-mapping strategies, and understanding evolutionary constraints on structural variation.

## 2 Results

### 2.1 Overview of the study

We developed MiXeR-SV, an extension of the established MiXeR framework[22–27], to quantify SV contributions to complex trait heritability using GWAS summary statistics (Figure 1). Our approach integrates three key components: (1) comprehensive ancestry-specific unified SNP-SV reference panels derived from the 1KGP, including whole-genome short-read sequencing of 3,202 samples[19] and 1,019 ONT long-read sequencing samples[20], establishing variant catalogs for EUR (*N* = 145 unrelated individuals) and EAS (*N* = 176) populations (Figure 1A); (2) LD score computation quantifying genome-wide correlation structure between SNPs and between SNPs and SVs (Figure 1B); and (3) a statistical framework based on GSA-MiXeR[25] that partitions heritability into SNP and SV contributions (Figure 1C). Our analysis pipeline tests for heritability enrichment in SVs relative to SNPs using likelihood ratio tests (LRT), followed by Wald tests to quantify fold-enrichment and assess whether it significantly exceeds unity (see Methods).

**Figure 1:**
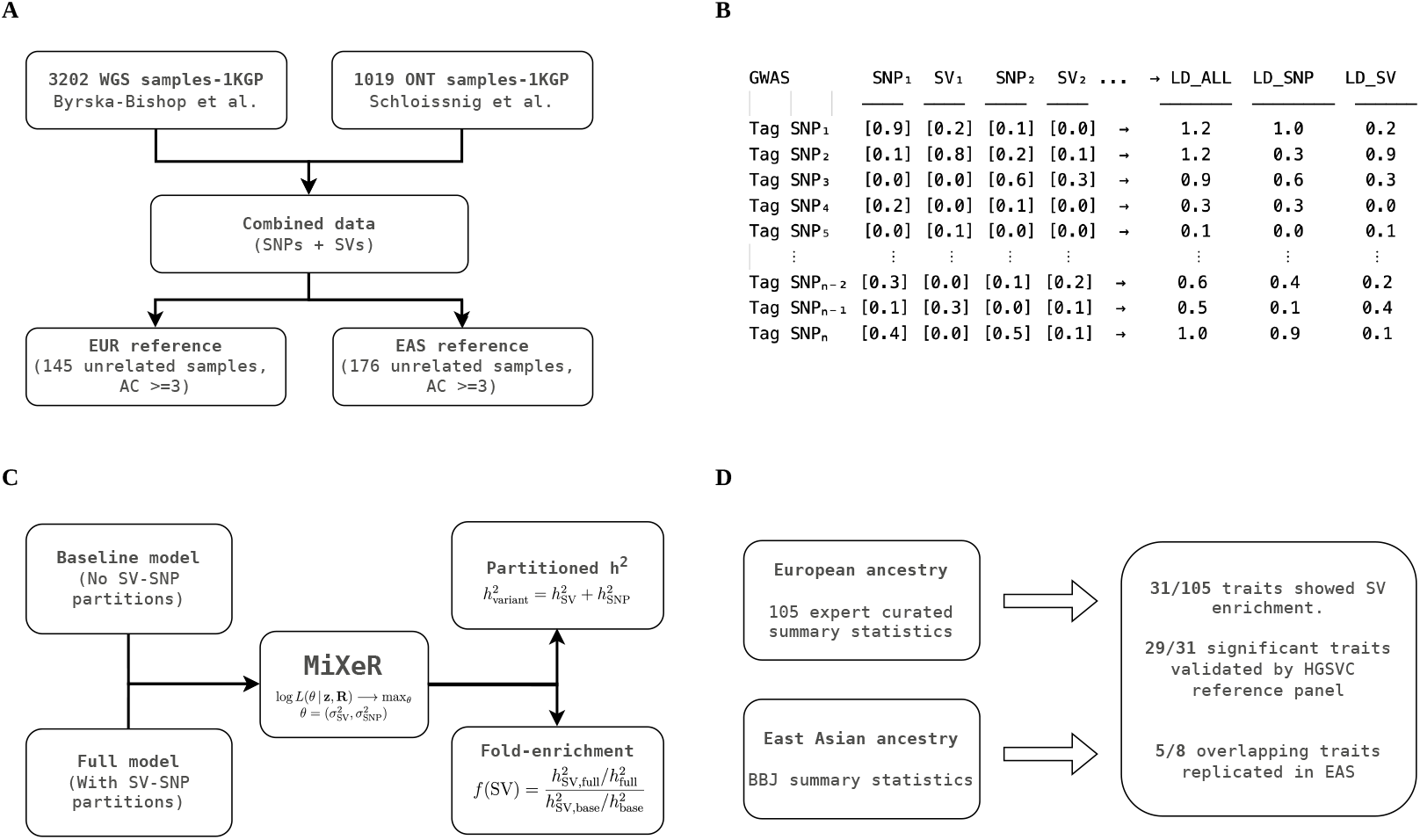
MiXeR-SV framework for quantifying structural variant heritability enrichment. **(A)** Ancestry-specific SV reference panels constructed from 3,202 whole-genome short-read sequences and 1,019 ONT long-read samples from the 1KGP, establishing unified SNP-SV catalogs for EUR (*N* = 145 unrelated individuals) and EAS (*N* = 176) populations (allele count ≥3). **(B)** LD score computation quantifies genome-wide correlation structure: LD_SV captures SNP-to-SV and LD_SNP captures SNP-to-SNP linkage patterns. **(C)** Genomic annotation stratified analysis based on GSA-MiXeR framework compares baseline model (no SV partitioning) versus full model (separate variant effect size variances into SNP and SV components) using LRT for significance and Wald test for fold-enrichment estimation. **(D)** Discovery analysis of 105 EUR traits identifies 31 with significant SV enrichment. Validation using an independent 1KGP HGSVC reference panel (EUR, *N* = 489) confirms enrichment consistency, and replication in the BBJ EAS cohort confirms five of eight overlapping traits.

We applied MiXeR-SV to 105 complex traits with publicly available GWAS summary statistics curated by Gazal et al.[28], combining diverse sources (UKBB, FinnGen, and many large-scale meta-analyses) and spanning neuropsychiatric, cardiometabolic, immune/autoimmune, cancer/aging, hematological, anthropometric, and metabolic/biomarker phenotypes. The discovery cohort comprised predominantly European ancestry samples ranging from approximately 10,000 to 2.5 million individuals (Supplementary Table S1). We identified 31 traits with genome-wide significant SV enrichment. To validate the robustness of our enrichment estimates independent of the ONT-based reference panel, we constructed a complementary SNP-SV reference panel leveraging the HGSVC phase 3 resource[21], which predict both SNPs and SVs from short-read sequencing data using a genome graph constructed from 214 human haplotypes anchored to a telomere-to-telomere (T2T) backbone. Applying for all 3,202 1KGP samples, we filtered to 489 unrelated EUR individuals, a sample size comparable to standard LDSC reference panels[16], and confirmed consistent SV heritability enrichment patterns across ancestries. We then analyzed eight overlapping traits from BBJ, with effective sample size ranging from 4,831 to 165,056 of EAS ancestry (Supplementary Table S2), and confirmed significant SV enrichment in five traits, further establishing the robustness and generalizability of our findings across diverse genetic backgrounds.

### 2.2 Common structural variants are well tagged by SNPs, enabling their effects to be detected through GWAS

A fundamental prerequisite for detecting SV effects through SNP-based GWAS is that SVs exhibit sufficient LD with genotyped or imputed SNPs. To assess this empirically, we analyzed genome-wide LD patterns between SNPs and SVs in EUR (*n* = 47, 991 SVs) and EAS (*n* = 47, 155 SVs) populations, stratified by variant type and MAF (Figure 2; Supplementary Figure 1). Insertions (INS) showed a slightly higher frequency for rare alleles, while deletions (DEL) and complex rearrangements (COM) exhibited frequency distributions comparable to SNPs (Figure 2A, Supplementary Figure 1A).

**Figure 2:**
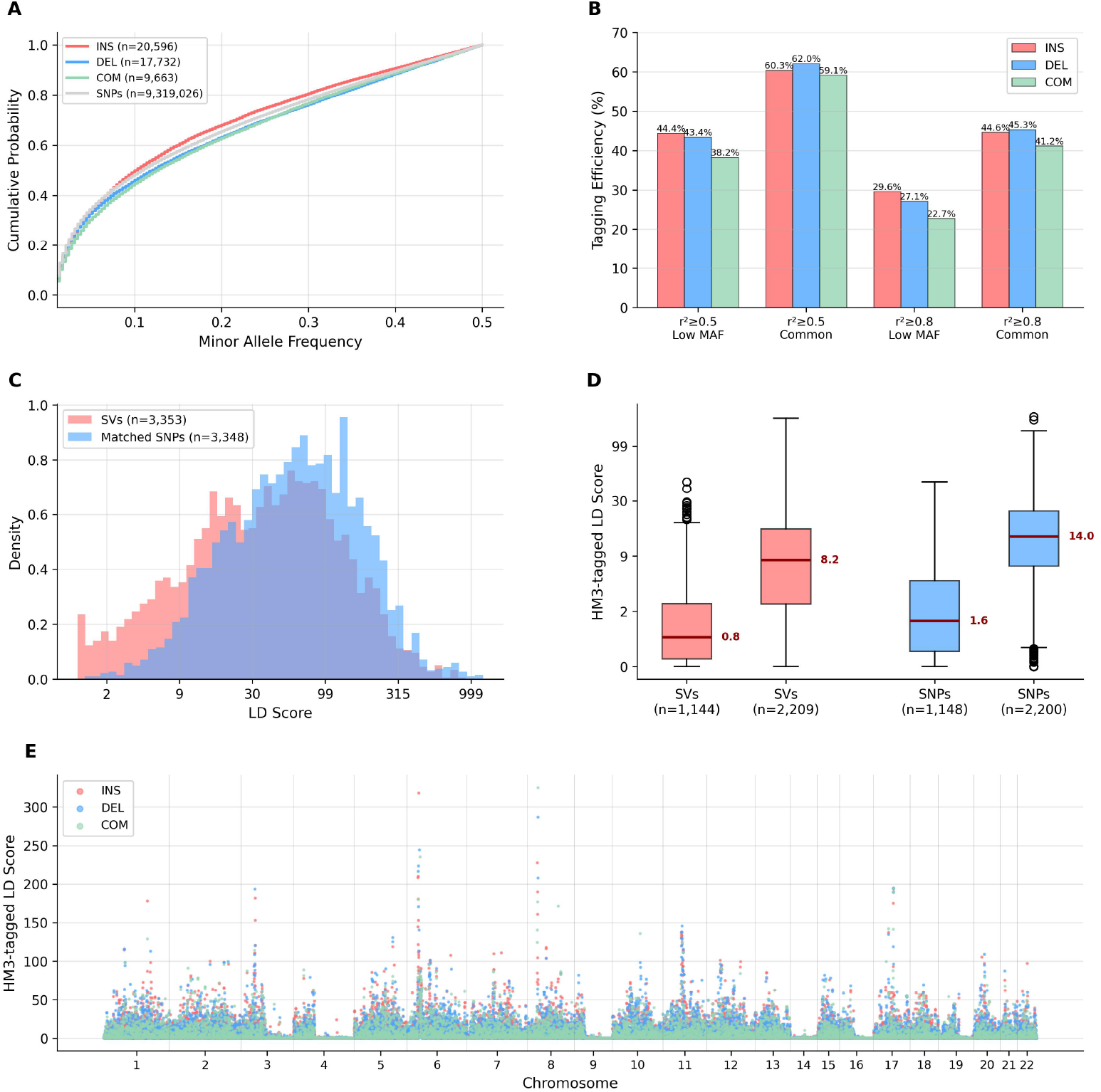
Structural variants exhibit substantial SNP tagging in European ancestry. **(A)** Cumulative MAF distributions show insertions have higher frequency for rare alleles, while deletions and complex rearrangements resemble SNPs (*N* = 47,991 SVs: 20,596 insertions, 17,732 deletions, 9,663 complex rearrangements). **(B)** SV tagging efficiency using all reference-panel SNPs as potential tags, stratified by MAF and *r*^2^ threshold. Common SVs (MAF≥0.05) show 59.1 to 62.0% tagging at *r*^2^ ≥ 0.5, whereas low-MAF SVs show 38.2 to 44.4%. At the stringent threshold (*r*^2^ ≥ 0.8), common SVs show 41.2 to 45.3% tagging and low-MAF SVs 22.7 to 29.6%. **(C)** Reference LD score density distributions on chromosome 1 (SVs *n* = 3,353, MAF-matched SNPs *n* = 3,348) using all SNPs as tags, showing that SVs have somewhat lower but overlapping LD scores compared with SNPs. **(D)** HM3-tagged LD scores on chromosome 1 after restricting tag variants to HapMap3 SNPs, stratified by MAF (SVs low MAF *n* = 1,144, SVs common *n* = 2,209, SNPs low MAF *n* = 1,148, SNPs common *n* = 2,200). **(E)** Genome-wide HM3-tagged LD scores for all 47,991 SVs reveal LD hotspots and inter-chromosomal heterogeneity in SV tagging by HapMap3 SNPs.

We first assessed tagging using all reference panel SNPs as potential tags. In EUR, 59.1 to 62.0% of common SVs (MAF≥0.05) had *r*^2^ ≥ 0.5 with at least one reference SNP, versus 38.2 to 44.4% of low-MAF SVs (Figure 2B). At the stringent threshold (*r*^2^ ≥ 0.8), 41.2 to 45.3% of common SVs showed very strong tagging, dropping to 22.7 to 29.6% for low-MAF SVs. MAF-matched LD score density distributions on chromosome 1 (*n* = 3, 353 SVs vs. *n* = 3, 348 SNPs) confirmed that SVs exhibit lower but substantially overlapping LD scores relative to matched SNPs overall (Figure 2C). EAS showed broadly consistent patterns: 57.7 to 61.3% of common SVs tagged at *r*^2^ ≥ 0.5, with lower low-MAF tagging (29.6 to 36.0%) than EUR, and similar patterns for chromosome 1 LD density (Supplementary Figure 1B,C).

We then confirmed these results using the more stringent criterion of HapMap3 (HM3) SNPs as tag variants only, which directly reflects the GWAS variants often used in downstream heritability analyses. HM3-tagged LD scores on chromosome 1 showed that common SVs retained substantial accessibility, in EUR, common SVs had a median HM3-tagged LD score of 8.2 versus 14.0 for matched common SNPs, while low-MAF SVs showed medians of 0.9 versus 1.6, respectively (Figure 2D). EAS exhibited similar patterns, with common SVs at 7.2 versus 13.3 for matched SNPs, and low-MAF SVs at 0.3 versus 0.9 (Supplementary Figure 1D). The reduction in absolute LD scores relative to the all-SNP analysis is expected given the sparser HM3 tag set, yet common SVs retain substantial LD scores, confirming that SV signals remain detectable through HM3-based GWAS summary statistics.

Genome-wide analysis of HM3-tagged LD scores revealed pronounced spatial heterogeneity, with consistent hotspot patterns across EUR and EAS (Figure 2E; Supplementary Figure 1E). The most prominent LD hotspots occurred at chromosomes 6, 8, and 11, where SVs exhibited exceptionally high LD scores reflecting extended haplotype structure. Several other genomic regions showed relatively low SV tagging density, indicating heterogeneous accessibility of SVs through SNP-based GWAS depending on local LD architecture. Despite differences in peak magnitudes driven by population-specific LD patterns and demographic histories, the spatial distribution of SV LD scores was remarkably consistent between EUR and EAS, suggesting conserved SV architecture and shared selective pressures across human populations.

Collectively, these analyses establish that common SVs are substantially accessible through existing SNP-based GWAS via LD tagging. Further confirmation using HM3-only tags demonstrates that this accessibility is preserved under the stringent reference used in our heritability framework. However, the incomplete tagging, particularly for low-MAF SVs and in genomic regions with poor LD coverage, indicates that SNP-based GWAS capture only a subset of total SV heritability contributions. Importantly, the strong concordance of tagging patterns between EUR and EAS populations supports the generalizability across ancestries and suggests that many GWAS signals currently attributed to SNPs may reflect underlying SV effects. This interpretation is further supported by recent evidence that common SVs can be accurately imputed from SNP array data using multi-ancestry long-read sequencing reference panels[15], confirming that SV information is recoverable from SNP-based data through LD-aware approaches.

### 2.3 Structural variants contribute substantially to complex trait heritability

Having established that common SVs are accessible through SNP-based GWAS, we applied MiXeR-SV (see Methods) to quantify SV heritability enrichment across 105 complex traits with publicly available summary statistics and identified 31 traits (29.5%) with significant enrichment of SV heritability(Bonferroni (BF)-corrected *P <* 4.8 × 10^−4^; Figure 3A,B; Supplementary Figure 2A; Supplementary Table S3). SV enrichment ranged from 6.6-fold for RBC count to 61.5-fold for type 1 diabetes, with median enrichment of 14.8-fold among significant traits. The heritability explained by SVs, denoted 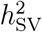, accounted for an average of 10.1% of total 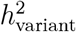 in significantly enriched traits (range 3.4 to 32.0%; Figure 3A), despite SVs comprising only approximately 0.6% of the total variants analyzed. The distribution of enrichment values showed a long tail extending beyond 40-fold (Supplementary Figure 2B, median across all 105 traits = 8.99-fold), with the most extreme values concentrated in immune-mediated and cancer phenotypes.

**Figure 3:**
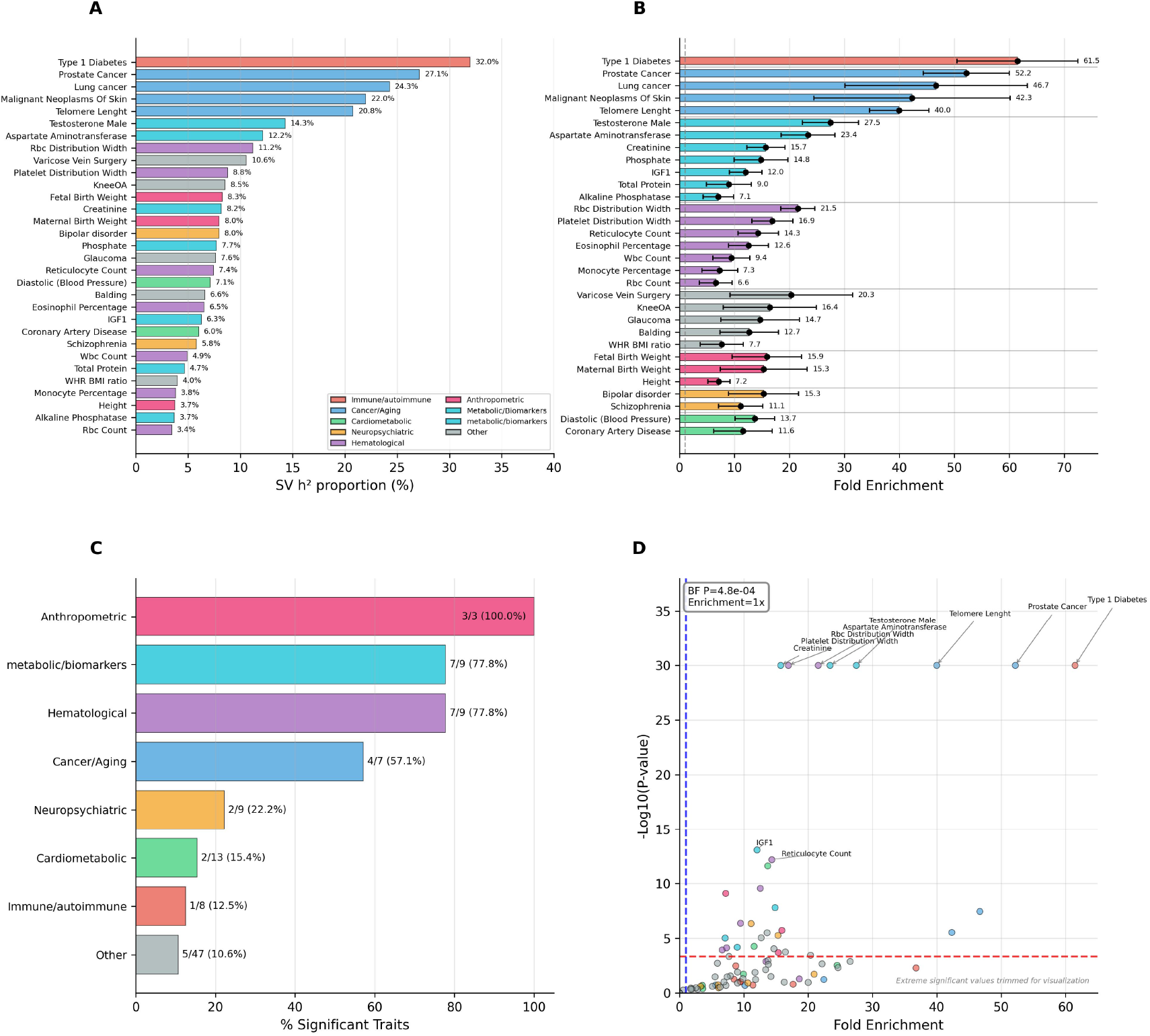
Structural variants show extensive heritability enrichment across complex traits. **(A)** Heritability partitioning for 31 traits with significant SV enrichment, showing the percentage of total variant heritability 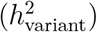 explained by SVs 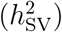, sorted by fold enrichment. Bars colored by trait category. **(B)** Forest plot displaying fold enrichment with 95% confidence intervals for all 31 significant traits, grouped by category and sorted by enrichment within each group. Gray horizontal lines separate trait categories; dashed vertical line indicates the null hypothesis (enrichment = 1). **(C)** Category-level enrichment summary showing the proportion of traits with significant SV enrichment within each of eight trait categories, sorted by percentage. Labels indicate the number of significant traits over total traits tested. **(D)** Volcano plot of fold enrichment versus statistical significance across all 105 analyzed traits. Points colored by trait category; horizontal red dashed line marks Bonferroni-corrected significance threshold (*P <* 4.8 × 10^−4^); vertical blue dashed line indicates null enrichment (enrichment = 1). Extreme significant values are trimmed at − log_10_(*P*) = 30 for visualization. Top 10 most significantly enriched traits are annotated.

Trait categories exhibited markedly heterogeneous patterns of enrichment (Figure 3C). Anthropometric traits showed enrichment in all analyzed traits (3/3 significant), though with modest effect sizes reflecting their highly polygenic architecture. Hematological and metabolic/biomarker traits demonstrated equally high categorical enrichment, each with 7 of 9 traits showing significant effects. Among hematological traits, RBC distribution width exhibited 21.5-fold enrichment, platelet distribution width 16.9-fold, and reticulocyte count 14.3-fold. In metabolic/biomarker traits, testosterone levels in males displayed 27.5-fold enrichment and aspartate aminotransferase 23.4-fold enrichment. Cancer and aging traits showed enrichment in 4 of 7 traits, led by prostate cancer with 52.2-fold enrichment and telomere length with 40.0-fold enrichment. In contrast, immune-mediated traits exhibited significant enrichment in only 1 of 8 traits, though type 1 diabetes showed the strongest enrichment genome-wide (61.5-fold).

Among neuropsychiatric traits, schizophrenia displayed 11.1-fold enrichment and bipolar disorder 15.3-fold enrichment, whereas other psychiatric traits including major depressive disorder showed no significant enrichment. Cardiometabolic traits exhibited selective enrichment in 2 of 13 traits, with diastolic blood pressure (13.7-fold) and coronary artery disease (11.6-fold) reaching significance while glycemic and lipid traits did not. Volcano plot analysis across all 105 traits confirmed that significant enrichment signals were concentrated in cancer, hematological, and metabolic domains (Figure 3D), with several traits exceeding 40-fold enrichment and maintaining genome-wide significance.

Partitioning absolute versus relative SV heritability contributions revealed distinct scenarios (Supplementary Figure 2D). Type 1 diabetes exhibited the highest relative contribution (32.0% of total heritability) despite modest absolute heritability 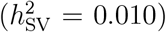, consistent with a small number of high-impact variants in the major histocompatibility complex region. Conversely, anthropometric traits such as height showed higher absolute SV heritability 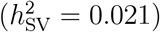 but a much lower relative contribution (3.7%), reflecting their highly polygenic backgrounds. Prostate cancer occupied a distinct position with both high absolute 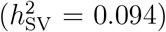 and high relative (27.1%) SV heritability contribution, suggesting SVs act as major contributors in a moderately heritable trait. This patterns demonstrates varying SVs contributions across diverse genetic architectures, from low-polygenic immune traits to highly polygenic anthropometric phenotypes. Finally, total variant heritability and SV enrichment estimates were largely independent from each other (Pearson *r* = 0.029, *P* = 7.7 × 10^−1^; Supplementary Figure 2C), indicating that enrichment patterns reflect trait-specific genetic architecture rather than total heritability.

These findings show that SVs contribute disproportionately to specific trait domains, particularly those involving hematopoiesis, immune function, and metabolic regulation. The observation that SV enrichment is independent of total heritability suggests biological rather than statistical origins, potentially reflecting regulatory hotspots where SVs modulate gene expression programs critical to trait manifestation [5, 29].

### 2.4 Human Genome Structural Variation Consortium reference panel validates SV enrichment findings

Given the limited sample size of the 1KGP ONT long-read cohort, we sought to validate the robustness of our enrichment estimates using a complementary reference panel that leverages the full 1KGP sample size. We constructed a second unified SNP-SV reference panel based on the HGSVC phase 3 resource[21] comprehensive variant call sets based on a genome graph built from 214 human haplotypes. Applying an identical quality control pipeline (Methods section), we obtained a reference panel of 489 unrelated EUR individuals, comparable in size to standard LDSC reference panels.

Using this new reference panel, 44 of 105 traits show significant SV enrichment (BF-corrected *P <* 4.8 × 10^−4^; Supplementary Table S4). Of the 31 ONT-significant traits, 29 (93.5%) were confirmed as significant by the HGSVC reference (Figure 4A), demonstrating near-complete replication of the primary findings. Enrichment estimates across all 105 traits were correlated between the two reference panels (Pearson *r* = 0.546, *P* = 1.8 × 10^−9^; Figure 4B), with substantially stronger agreement among the 29 traits significant under both references (Pearson *r* = 0.745, *P* = 3.6 × 10^−6^; Figure 4C). Mean enrichment estimates were slightly higher under the ONT reference (19.7 *±* 14.6-fold) than HGSVC (17.0 *±* 12.5-fold), consistent with the ONT catalog capturing a broader SV landscape including difficult-to-detect insertions and complex rearrangements that may not be fully represented in the graph-based HGSVC reference.

**Figure 4:**
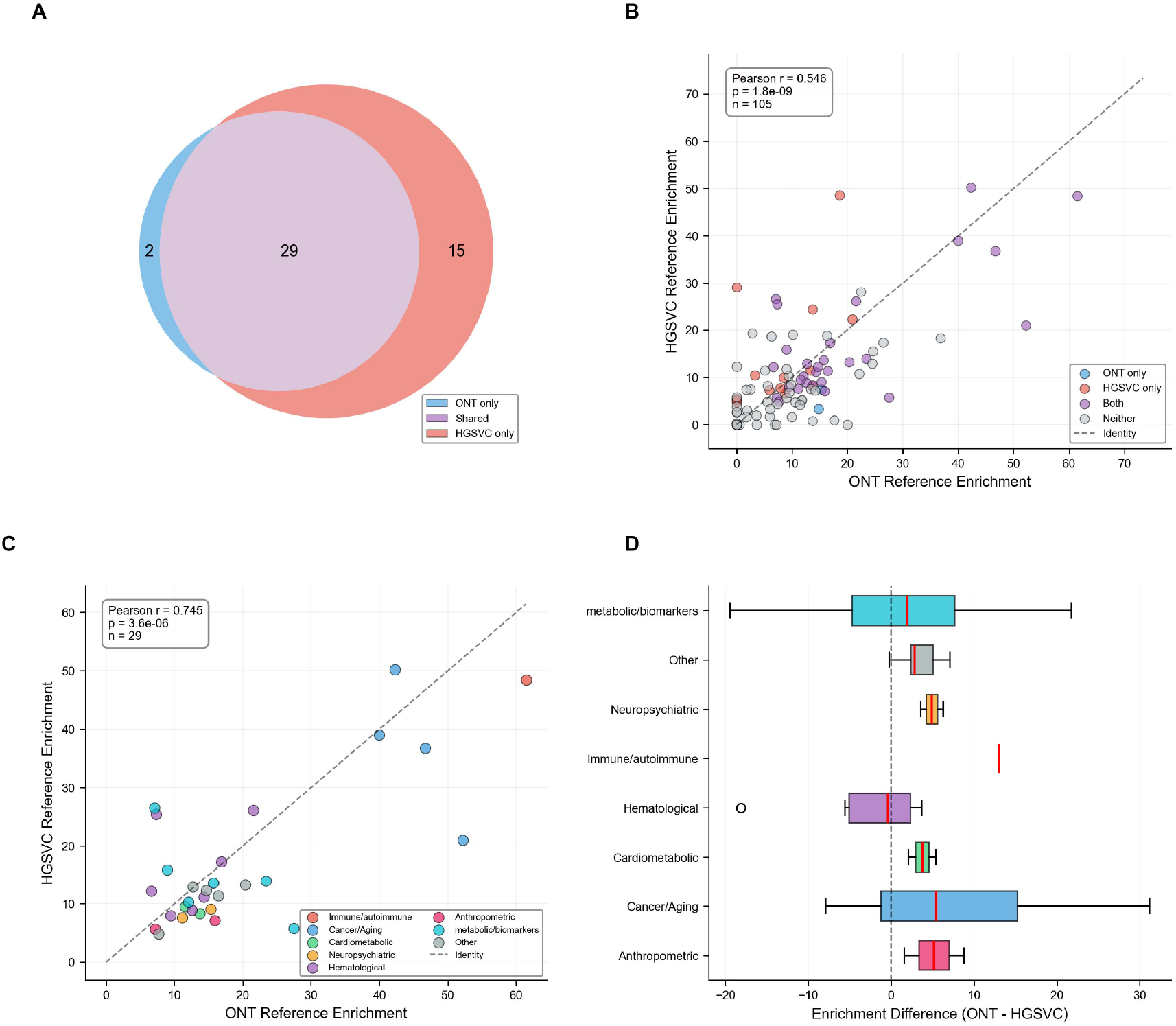
Complementary HGSVC reference panel validates SV heritability enrichment. **(A)** Venn diagram showing overlap of significantly enriched traits between the primary ONT-based reference (31 traits) and the complementary HGSVC-based reference (44 traits). Of 31 ONT-significant traits, 29 (93.5%) are confirmed by HGSVC; 15 additional traits reach significance exclusively under HGSVC. **(B)** Scatter plot of fold enrichment estimates for all 105 traits under both reference panels (Pearson *r* = 0.546, *P* = 1.8 × 10^−9^, *n* = 105), colored by significance category (ONT only, HGSVC only, both, or neither). Dashed line indicates identity (*y* = *x*). **(C)** Scatter plot restricted to the 29 traits significant under both reference panels (Pearson *r* = 0.745, *P* = 3.6 × 10^−6^, *n* = 29), colored by trait category. **(D)** Horizontal boxplot of per-category enrichment differences (ONT − HGSVC) among shared significant traits.

The two ONT-significant traits that did not reach significance under HGSVC, phosphate (14.8-fold, ONT) and maternal birth weight (15.3-fold, ONT), both showed directionally consistent enrichment estimates under HGSVC, suggesting insufficient power rather than discordant biological signal. Conversely, the 15 traits that reached significance exclusively under HGSVC but not ONT included notable findings such as clonal hematopoiesis of indeterminate potential (CHIP; 48.6-fold), Alzheimer’s disease (22.2-fold), varicose veins (24.3-fold), and venous thromboembolism (29.0-fold), which may reflect additional statistical power afforded by the larger HGSVC reference sample size. Per-category enrichment differences between ONT and HGSVC were modest for most trait domains, with the largest variability observed in cancer/aging and metabolic/biomarker traits (Figure 4D), likely reflecting differential representation of specific SVs between the graph-based imputed and long-read directly called catalogs.

Collectively, the 93.5% concordance of significantly enriched traits and the strong correlation of SV enrichment estimates among shared traits establish that the primary MiXeR-SV findings are robust to the choice of reference panel. The additional traits reaching significance under HGSVC suggest that the ONT-based analysis may represent a conservative estimate of the total number of traits with genuine SV heritability enrichment, and that expanding the ONT long-read reference to population scale could further increase discovery power.

### 2.5 SVs contribute distinct heritability signal consistent with the missing heritability gap

To assess whether SVs capture genuinely distinct heritability signal beyond standard SNP-based models, we compared total *h*^2^ estimates between the full MiXeR-SV model and a constrained model in which the SV-specific effect size variance is fixed to zero (Supplementary Table S5). Among the 31 traits with significant SV enrichment, the full model consistently yielded higher total *h*^2^ estimates, with a median ratio of 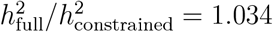 (mean = 1.049, range 1.013 to 1.185; Figure 5A). The magnitude of this gain scaled directly with SV enrichment (Pearson *r* = 0.989, *P* = 8.9 × 10^−26^, *n* = 31), indicating that the additional heritability captured by the full model reflects genuine SV signal proportional to enrichment strength rather than a statistical artifact of added model complexity. Eight traits showed *h*^2^ gains exceeding 5%, and five exceeded 10%, with the largest improvements in type 1 diabetes (+18.5%; enrichment 61.5×) and prostate cancer (+17.2%; enrichment 52.2×). Traits lacking significant enrichment showed negligible differences (*h*^2^ ratio ≈ 1.00), confirming that the full model does not artificially inflate heritability estimates in the absence of genuine SV signal.

**Figure 5:**
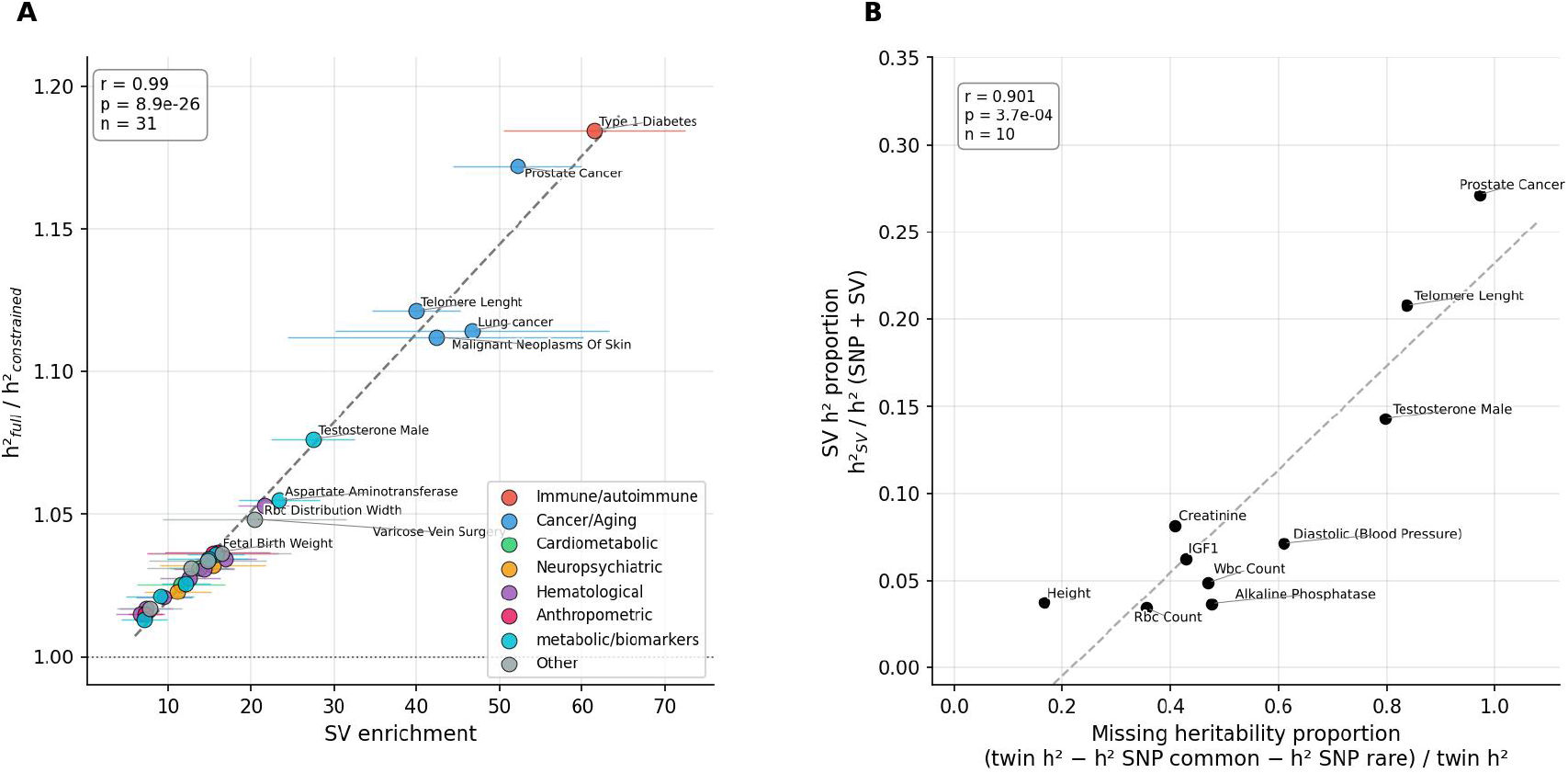
SVs contribute distinct heritability signal consistent with the missing heritability gap. **(A)** Scatter plot of SV fold enrichment versus the ratio 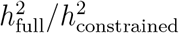 for 31 traits with significant SV enrichment (Supplementary Table S5). The full model estimates a separate SV-specific effect size variance; the constrained model fixes this variance to zero. Points are coloured by trait category; error bars represent 95% confidence intervals. The dashed line shows the linear regression fit (Pearson *r* = 0.989, *P* = 8.9 × 10^−26^, *n* = 31); the dotted horizontal line indicates a ratio of 1.0. **(B)** Scatter plot of missing heritability proportion against SV heritability proportion for 10 traits with available twin heritability estimates. Missing heritability is defined as 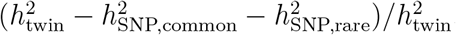, where total SNP-based heritability is from Wainschtein et al. [3] and twin estimates are from published classical twin studies (Supplementary Table S6). The dashed line shows the linear regression fit (Pearson *r* = 0.901, *P* = 3.7 × 10^−4^, *n* = 10).

To contextualize SV contributions within the broader heritability landscape, we compared SV heritability fractions against the missing heritability of each trait, defined as the proportion of twin-based heritability not explained by common and rare SNPs combined. For the ten significantly enriched traits at the intersection of our results, those reported by Wainschtein et al.[3], and those with available twin heritability estimates, we obtained total SNP-based heritability (common plus rare variants) from Wainschtein et al.[3] and twin heritability estimates from published classical twin studies (Supplementary Table S6). We defined missing heritability to twin heritability proportion (distinct from the family-based missing heritability defined in[3]) as 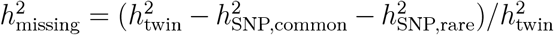. As shown in Figure 5B, the proportion of total heritability explained by SVs was strongly correlated with the missing heritability fraction across these ten traits (Pearson *r* = 0.901, *P* = 3.7 × 10^−4^, *n* = 10). Traits with the largest missing heritability, including prostate cancer and telomere length, also showed the highest SV heritability proportions (27% and 21%, respectively). Conversely, height, for which SNPs already account for most twin heritability, showed the most modest SV contribution (3.7%). Together, these findings are consistent with SVs contributing distinct heritability signal that is systematically underestimated when SV architecture is ignored in partitioned heritability models, and with SVs being under-tagged by common variant arrays rather than truly absent from the genetic architecture of complex traits.

### 2.6 Cross-ancestry replication in Biobank Japan confirms SV enrichment

To validate cross-ancestry generalizability of SV enrichment patterns and assess whether our findings reflect shared biology versus population-specific LD structure, we tested enrichment in eight traits from BBJ that overlapped with significant traits identified in the discovery phase (EAS ancestry, *N* ≤ 165, 056). We required both statistical significance (LRT and Wald test, *P <* 0.05) similar to the primary analysis (Supplementary Table S7).

Five of tested traits (62.5%) showed robust replication with consistent enrichment estimates across ancestries (Figure 3A). Blood cell traits demonstrated strong replication: red blood cell count showed 6.6-fold enrichment in EUR versus 16.2-fold in BBJ (Wald *Z* = 3.09, LRT *χ*^2^ = 27.9), and white blood cell count showed 9.4-fold versus 16.5-fold enrichment (Wald *Z* = 3.03, LRT *χ*^2^ = 25.9). Height replicated with 7.2-fold enrichment in EUR versus 10.2-fold in BBJ (Wald *Z* = 3.77, LRT *χ*^2^ = 130.2). Prostate cancer enrichment was highly consistent across ancestries (EUR: 52.2×, BBJ: 55.8×; Wald *Z* = 4.05, LRT *χ*^2^ = 10.2). Diastolic blood pressure showed moderate enrichment in EUR (13.7×) and higher enrichment in BBJ (25.6×; Wald *Z* = 2.74, LRT *χ*^2^ = 7.3).

Among successfully replicated traits, enrichment estimates were highly correlated across ancestries (Pearson *r* = 0.982, *P* = 2.9 × 10^−3^, *N* = 5; Figure 3B), demonstrating that SV contributions to complex traits are largely shared between EUR and EAS populations. The strong correlation and overlapping confidence intervals between EUR and BBJ estimates for all five replicated traits indicate that the observed enrichment reflects genuine biological signal rather than population-specific confounding from LD structure or demographic history.

Three traits failed to replicate, likely reflecting insufficient statistical power or true population-specific genetic architecture. Type 1 diabetes, despite showing the strongest enrichment in EUR discovery (61.5×), exhibited a nominally consistent but statistically non-significant estimate in BBJ (32.0 *±* 45.2×; Wald *Z* = 0.69, LRT *χ*^2^ = 0.16), where the large standard error reflects insufficient statistical power given the small BBJ effective sample size (*N*_eff_ = 4,831). Lung cancer failed LRT significance in BBJ (LRT *χ*^2^ = 3.79, *P >* 0.05) despite a nominally large point estimate (174.6 *±* 42.1×; Wald *Z* = 4.12), suggesting that the BBJ lung cancer sample size (*N*_eff_ = 17,334) is insufficient for reliable model fitting. Glaucoma showed no detectable enrichment in BBJ (0×; LRT *χ*^2^ = 0.24), consistent with established population differences in glaucoma genetic architecture, where primary angle-closure glaucoma predominates in EAS populations while primary open-angle glaucoma is more common in EUR[30, 31].

Collectively, the 62.5% replication rate and high correlation among successfully powered traits establish that SV enrichment patterns are robust across ancestries for traits with shared genetic architecture, while highlighting expected population specificity for traits with distinct biological subtypes. These findings validate MiXeR-SV’s ability to detect genuine SV effects and demonstrate that SV contributions to blood cell phenotypes, anthropometric traits, and prostate cancer represent conserved features of human genetic architecture.

## 3 Discussion

Using recent long-read sequencing reference data, we developed MiXeR-SV to quantify SV contributions to complex trait heritability using GWAS summary statistics. Applying this framework to 105 complex traits, we identified 31 traits with significant SV enrichment, with fold-enrichments varying greatly across trait categories. Robustness of the primary findings was confirmed using a complementary HGSVC-based reference panel, which replicated the majority of ONT-significant traits and revealed additional signals enabled by the larger reference sample size. The utility of our approach is supported by the increasing accessibility of high-quality SV reference catalogs. ONT long-read sequencing now enables sequence-resolved SV discovery at population scale [20], while pangenome-based SV genotyping approaches provide complementary and scalable alternatives [21, 32]. Cross-ancestry replication in BBJ further confirmed enrichment in the majority of tested traits with near-perfect correlation between EUR and EAS estimates. Together, these findings establish that SVs contribute substantially to specific complex trait domains and that these contributions can be quantified from existing SNP-based data without requiring GWAS data for SVs.

The heterogeneous enrichment patterns across complex trait categories reveal distinct patterns of genetic architectures. Hematological and metabolic/biomarker traits showed the highest categorical enrichment rates, with SV explaining a disproportionate share of total heritability, indicating systematic SV effects on blood cell phenotypes and metabolic regulation. In contrast, immune-mediated traits showed enrichment in only one trait, though type 1 diabetes exhibited the strongest genome-wide signal by a wide margin. Anthropometric traits showed consistent but lower SV enrichment, suggesting that SV contributions to height and birth weight follow a broadly polygenic architecture similar to that described for SNPs, rather than concentrated regulatory effects.

The trait-specific enrichment patterns we observe align with functional genomic evidence demonstrating that SVs exert disproportionate effects on gene regulation[9]. eQTL mapping in human populations reveals that common SVs are causal at 2.66% of eQTLs despite comprising only 0.25% of variants, representing a 10.5-fold enrichment[33]. This enrichment is even more pronounced for splicing QTLs, with a recent cattle long-read sequencing study identifying up to 12-fold enrichment for SVs when accounting for LD[8]. The mechanistic basis for these effects involves large-scale disruption of three-dimensional chromatin architecture. SVs that alter topologically associating domain (TAD) boundaries cause pathogenic rewiring of gene-enhancer interactions through “enhancer hijacking”, whereby removal or displacement of TAD boundaries exposes promoters to ectopic regulatory elements[34]. Notably, SV-eQTLs affect an average of 1.82 nearby genes compared to 1.09 genes for SNV-eQTLs[33], providing a mechanism by which individual SVs may exert pleiotropic effects on complex traits through coordinated dysregulation of multi-gene regulatory domains.

Our observation of extensive enrichment in hematological traits is consistent with the critical role of SVs in blood diseases. A recent whole-genome sequencing study in 50,675 individuals identified 21 independent SVs associated with quantitative blood cell traits[35]. Classical examples include alpha-thalassemia, where large deletions removing HBA1 and HBA2 on chromosome 16p13.3 cause microcytic hypochromic anemia[36], and BCL11A deletions that result in persistent fetal hemoglobin expression[37]. These Mendelian deletion syndromes demonstrate that hematopoietic pathways are particularly vulnerable to dosage imbalance from structural variation. Similarly, the enrichment observed across anthropometric traits is compatible with known examples such as ACAN deletions causing familial short stature through haploinsufficiency[38]. For cancer phenotypes, the strong enrichment in prostate cancer and lung cancer is consistent with known germline copy number variant associations with cancer risk[39], and extreme enrichment for telomere length aligns with established roles for structural variants at telomere maintenance loci[40].

Cross-ancestry replication provides evidence for shared biological mechanisms underlying SV enrichment. The high replication rate, and consistent fold enrichments among successfully replicated traits establish that SV contributions to blood cell phenotypes, anthropometric traits, and prostate cancer are conserved between EUR and EAS populations. Replication failures in type 1 diabetes and lung cancer likely reflect insufficient statistical power in the smaller BBJ cohorts rather than true biological inconsistency. Non-replicating results for glaucoma may reflect population-specific genetic architecture, consistent withthe differing predominance of primary angle-closure and open-angle subtypes between EAS and EUR populations[30, 31], and suggests that MiXeR-SV captures genuine biological differences rather than methodological artifacts. The additional traits identified exclusively by the HGSVC reference panel, including clonal hematopoiesis, Alzheimer’s disease, and venous thromboembolism, suggest that the primary ONT-based analysis is conservative and that expanding long-read reference panels to population scale will increase discovery power. Furthermore, the strong correlation between SV enrichment and the twin-based missing heritability fraction suggests that SVs capture genuinely distinct genetic signal that escapes detection in conventional SNP-based analysis.

Our findings have immediate implications for polygenic risk prediction and GWAS interpretation. For traits with high SV enrichment, current polygenic risk prediction models relying on SNP tagging likely miss substantial predictive signal, as our analyses demonstrate that common SVs show incomplete LD coverage by HapMap3 SNPs, particularly in genomic regions with heterogeneous tagging density. Traits where SVs explain a substantial fraction of total heritability represent priority targets for incorporating direct SV genotypes from long-read sequencing. Fine-mapping efforts at enriched GWAS loci should prioritize comprehensive SV catalogs, as SNP-only credible sets may systematically exclude causal SVs in weak LD with genotyped markers.

The framework’s limitations include reliance on LD tagging, which captures only a subset of structural variation, and a primary reference panel of modest size from the 1KGP ONT cohort. Although the complementary HGSVC reference panel confirms the robustness of our findings, enrichment estimates represent lower bounds on true SV contributions. Future work incorporating population-scale long-read sequencing cohorts, expanded ancestry diversity, and integration with functional genomic data will enable identification of specific causal SVs and their mechanistic effects.

Overall, MiXeR-SV demonstrates that SVs make substantial, trait-specific contributions to heritability that are quantifiable from existing GWAS summary statistics. We establish a clear hierarchy of SV enrichment across 105 complex human traits and validate these patterns across ancestries. Finally, our work helps prioritize traits for future long-read and pangenome sequencing studies, guiding the deployment of these resource-intensive technologies toward settings where they are most likely to yield biological insight.

## 4 Methods

### 4.1 MiXeR statistical model

We extend the MiXeR framework[22, 23] to partition total variant heritability into SV and SNP components. Under an additive genetic model, a standardized (zero mean, unit variance) quantitative phenotype *y* ∈ ℝ can be modeled as

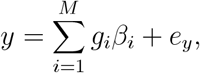

where *M* denotes the total number of genetic variants in the reference panel, *g*_*i*_ is the number of reference alleles (centered, but not variance standardized) for the *i*-th variant, *β*_*i*_ is the per-allele effect size, and 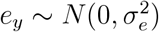 is the residual term capturing environmental effects and non-additive genetic contributions.

Specific prior distribution on effect sizes *β*_*i*_ encodes various genetic architectures of complex traits, for example 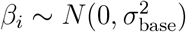 assumption is widely known as *infinitesimal model*. In MiXeR-SV we employ the *baseline model* (accounting for MAF-dependent architecture) and *full model* (additionally accounting for genomic regions of interest, such as SVs and SNPs). Let *A*_SV_ and *A*_SNP_ denote the SV and SNP variant sets in the reference panel.

#### The baseline model

is defined by 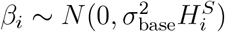, where the *S* parameter controls the effect size distribution with regards to allele frequency, similarly to LDAK and SumHer models. The original GCTA model assumes *S* = −1, so that each SNP contributes equally to heritability regardless of its allele frequency. In this notation, Var(*g*_*i*_) = *H*_*i*_ = 2*f*_*i*_(1 − *f*_*i*_), thus under the baseline model, total trait’s variant-based heritability 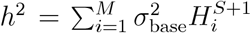. For any subset of variants *A*, the baseline-model heritability is 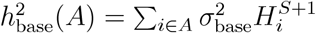.

#### The full model

extends the baseline model by allowing different effect-size variances for SVs and SNPs:

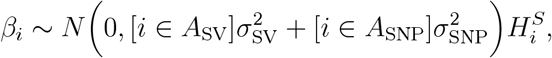

leading to 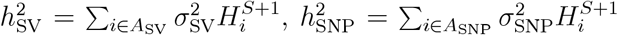, _and 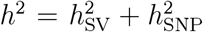_ estimates for SV, SNP, and total heritability, respectively. *Fold-enrichment of SV heritability* relative to the baseline model is then defined as

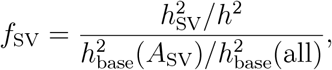

and *heritability proportions* within the full model are

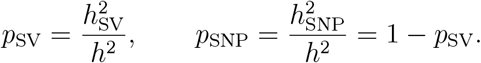

The implementation follows GSA-MiXeR[25], where model parameters are estimated by maximizing the likelihood of GWAS summary statistics (marginal SNP-trait associations) under the model, accounting for LD structure in the reference panel. Parameters are initialized using method-of-moments (MoM) estimates as starting points for maximum likelihood estimation (MLE) implemented via PyTorch `torch.autograd` library for automatic gradient computation in Adam optimization.

SEs for parameter estimates are derived from the observed Fisher information matrix, defined as the negative Hessian matrix of the log-likelihood function log *L*(*θ*|**z**) evaluated at the maximum likelihood estimate *θ*^∗^:

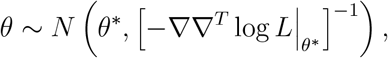

where 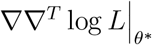 denotes the matrix of second partial derivatives with respect to model parameter *θ*, computed using `torch.autograd.functional.hessian`. For compound functions of multiple parameters, such as fold enrichment of heritability in a genomic region, we sample N=100 realizations of the parameters vector from the above posterior distribution, calculate the compound function for each realization of the parameter vector, and report the standard deviations across the realizations. If the Hessian matrix was not positive definite, we use marginal errors of fitted parameters.

Model fit was assessed using log-likelihoods log *L*_base_ and log *L*_full_, computed under each model for the observed z-score distribution. A likelihood ratio test (LRT) was used to determine whether inclusion of the additional parameter, 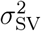, in the full model significantly improved the fit compared to the base model:

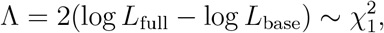

Significance was assessed at BF-corrected threshold *α* = 0.05*/*105 = 4.8 × 10^−4^ for 105 traits in the discovery analysis. To confirm that enrichment was in the positive direction (i.e., 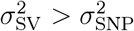), we performed a one-sided Wald test on the fold-enrichment, *Z*_wald_ = (*f*_SV_ − 1)*/*SE[*f*_SV_], where the standard error was obtained from the Fisher information matrix. A trait was classified as significantly enriched only if both the LRT and Wald test reached the BF threshold.

For BBJ validation, we applied MiXeR-SV framework using ancestry-matched reference panels (EAS, *N* = 176 unrelated individuals) to compute LD scores and estimate model parameters. Replication required both statistical tests (improved fit of the full model vs null model, and positive enrichment in SVs) to pass nominal significance thresholds (*P <* 0.05, without BF correction given the validation context. For correlation analysis, we excluded traits that either failed replication and focusing on robustly estimated effects to avoid outlier influence.

### 4.2 Reference panel construction

We constructed ancestry-specific unified SNP-SV reference panels from two complementary 1KGP resources, both processed under identical quality control criteria (biallelic variants only, Hardy-Weinberg equilibrium *P* ≥ 10^−6^, ≤ 5% missing genotypes, AC ≥ 3 per ancestry, unrelated individuals only[16], short indels excluded).

#### 4.2.1 ONT-based primary reference

We integrated phased SNP genotypes for 3,202 individuals from the 1KGP high-coverage short-read dataset (GRCh38, released 2020-10-28)[19] with phased SV genotypes from the 1KGP ONT long-read dataset[20], comprising 1,019 individuals (release v1.1, shapeit5-phased callset) with insertions, deletions, and complex rearrangements ranging from 50 bp to several megabases. SNP VCF files were downloaded for autosomes (chromosomes 1 to 22), restricted to biallelic SNPs, and annotated with dbSNP rsIDs (build 151) using BCFtools[41], retaining only SNPs with a valid rsID annotation. SNP and SV VCF files were merged using BCFtools concat to create unified variant catalogs. After quality control, this yielded 145 unrelated EUR and 176 unrelated EAS individuals.

#### 4.2.2 HGSVC-based complementary reference

We obtained variant genotypes for all 3,202 1KGP individuals from the HGSVC phase 3 resource[21], which provides genome-wide SNP and SV genotypes predicted by PanGenie[42] using a genome graph constructed from 214 human haplotypes anchored to a telomere-to-telomere backbone. We filtered the resulting genotype calls to retain only SNPs and SVs, excluding short indels. SNPs were annotated with dbSNP rsIDs (build 151) using BCFtools[41] and restricted to those with a valid rsID annotation, consistent with the ONT-based panel. After applying the same quality control pipeline, this yielded 489 unrelated EUR individuals.

#### 4.2.3 LD reference panel preparation

For both reference panels, genotype data were converted to PLINK binary format (.bed/.bim/.fam) using PLINK 2.0[43], with separate files for each chromosome. Pairwise LD values were precomputed in 2.5 Mb sliding windows using PLINK 1.9 (`–r2 –ld-window-kb 2500 –ld-window-r2 0.1`), storing all *r*^2^ ≥ 0.1 values for downstream LD score computation.

### 4.3 GWAS summary statistics

#### 4.3.1 European ancestry discovery cohort

EUR GWAS summary statistics were selected from a curated list of 107 independent traits developed by Legros, Kim, and Gazal (available at https://doi.org/10.5281/zenodo.10515792). This collection was designed to minimize sample overlap and genetic correlation, with traits defined as independent if they had either non-overlapping samples or genetic correlation 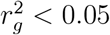 (for UK Biobank-derived traits). The curation process prioritized high-quality summary statistics through multiple filters: (1) starting from 231 summary statistics representing 63 independent traits from Gazal et al.[44], Global Biobank Meta-analysis Initiative, Pan-ancestry UKBB, and other public repositories; (2) selecting traits with heritability *Z*-scores *>* 6 (estimated via stratified LD score regression with the baseline-LD model[17]), with exceptions for immune-related traits that remained informative despite lower heritability; (3) prioritizing disease phenotypes over quantitative traits when genetic correlation indicated redundancy; and (4) removing traits with substantial sample overlap identified through cross-trait LD score regression intercepts [28]. Of the 107 curated traits, two were excluded from the analysis. One was unavailable due to data access restrictions, and one duplicated in the original curation.

The final 105 traits represent eight major phenotypic domains: neuropsychiatric disorders (schizophrenia, depression, ADHD, Alzheimer’s disease, Parkinson’s disease, autism spectrum disorder, anorexia nervosa, bipolar disorder, Tourette syndrome; *N* = 9 traits), cardiometabolic diseases (coronary artery disease, type 2 diabetes, atrial fibrillation, stroke, heart failure, abdominal aortic aneurysm, venous thromboembolism, BMI, blood pressure, cholesterol, HDL, LDL; *N* = 13 traits), immune/autoimmune conditions (asthma, inflammatory bowel disease, rheumatoid arthritis, type 1 diabetes, primary biliary cirrhosis, lupus, celiac disease; *N* = 8 traits), cancer and aging-related traits (prostate cancer, breast cancer, lung cancer, esophageal cancer, malignant neoplasms of skin, telomere length, parental lifespan; *N* = 7 traits), hematological traits (white blood cell count, red blood cell count, eosinophil percentage, red blood cell distribution width, reticulocyte count, platelet distribution width, monocyte percentage, basophil percentage, CHIP; *N* = 9 traits), anthropometric traits (height, fetal birth weight, maternal birth weight; *N* = 3 traits), metabolic biomarkers (total protein, creatinine, IGF1, aspartate aminotransferase, testosterone, alkaline phosphatase, phosphate, vitamin D, total bilirubin; *N* = 9 traits), and other traits including osteoarthritis, respiratory conditions, gastrointestinal disorders, lifestyle factors, and cognitive measures (*N* = 47 traits). Sample sizes ranged from 10,263 to 2,520,920 with a median of 408,112 individuals.

#### 4.3.2 East Asian ancestry validation cohort

For cross-ancestry validation, we analyzed eight BBJ traits [45] with phenotypic matches in our EUR discovery set, selected based on availability and trait overlap. These comprised: height (anthropometric; *N* = 165, 056), red blood cell count and white blood cell count (hematological; *N* = 153, 512 and 154, 355), diastolic blood pressure (cardiometabolic; *N* = 145, 515), glaucoma (*N* = 32, 182), prostate cancer and lung cancer (cancer/aging; *N* = 21, 263 and 17,334), and type 1 diabetes (immune/autoimmune; *N* = 4, 831). All BBJ summary statistics were derived from Japanese individuals and obtained from the BBJ public repository (see Data Availability). Sample sizes represent effective sample sizes for binary traits and total sample sizes for quantitative traits (Supplementary Table S2).

#### 4.3.3 Quality control and harmonization

All GWAS summary statistics were harmonized to rsIDs and GRCh37/hg19 coordinates with consistent allele coding and aligned to the reference LD panel. We excluded variants with strand ambiguities (A/T or G/C), MAF *<* 5%, low imputation quality (INFO *<* 0.8), missing effect estimates or SEs, non-autosomal variants, variants within the major histocompatibility complex region (chr6:25–34 Mb), duplicated SNPs, variants with extreme or low effective sample size, and variants not present in the reference LD panel.

## Ethics statement

This study used only publicly available data. All data were obtained from previously published studies or public repositories with existing ethical approvals.

## Data availability

All data used in this study are publicly available. The ONT long-read data were obtained from variant call set was obtained from https://ftp.1000genomes.ebi.ac.uk/vol1/ftp/data_collection.s/1KG_ONT_VIENNA/release/v1.1/, the high-coverage short-read data were obtained from https://ftp.1000genomes.ebi.ac.uk/vol1/ftp/data_collections/1000G_2504_high_coverage/working/20201028_3202_phased/, and the HGSVC variant call set was obtained from https://ftp.1000genomes.ebi.ac.uk/vol1/ftp/data_collections/HGSVC3/. GWAS summary statistics were obtained from https://doi.org/10.5281/zenodo.10515792 and https://pheweb.jp/downloads. The LD reference data generated for MiXeR-SV will be made publicly available upon completion of peer review.

## Code availability

Data and code for all analyses can be accessed at upon completion of peer review https://github.com/precimed/mixer_sv

## Author contributions

Conceptualization: DTN, OF, AAS, AMD, OAA; Data curation: DTN, OF, AAS, JF, NP, NSV; Formal analysis: DN, OF; Funding acquisition: OAA, AMD; Software: DTN, OF, AAS; Visualization: DTN, OF; Writing – original draft: DTN, OF; Writing – review & editing: all listed authors.

