## Supplementary Figure for "Structural variants contribute substantially to complex trait heritability"

### Supplementary Figures

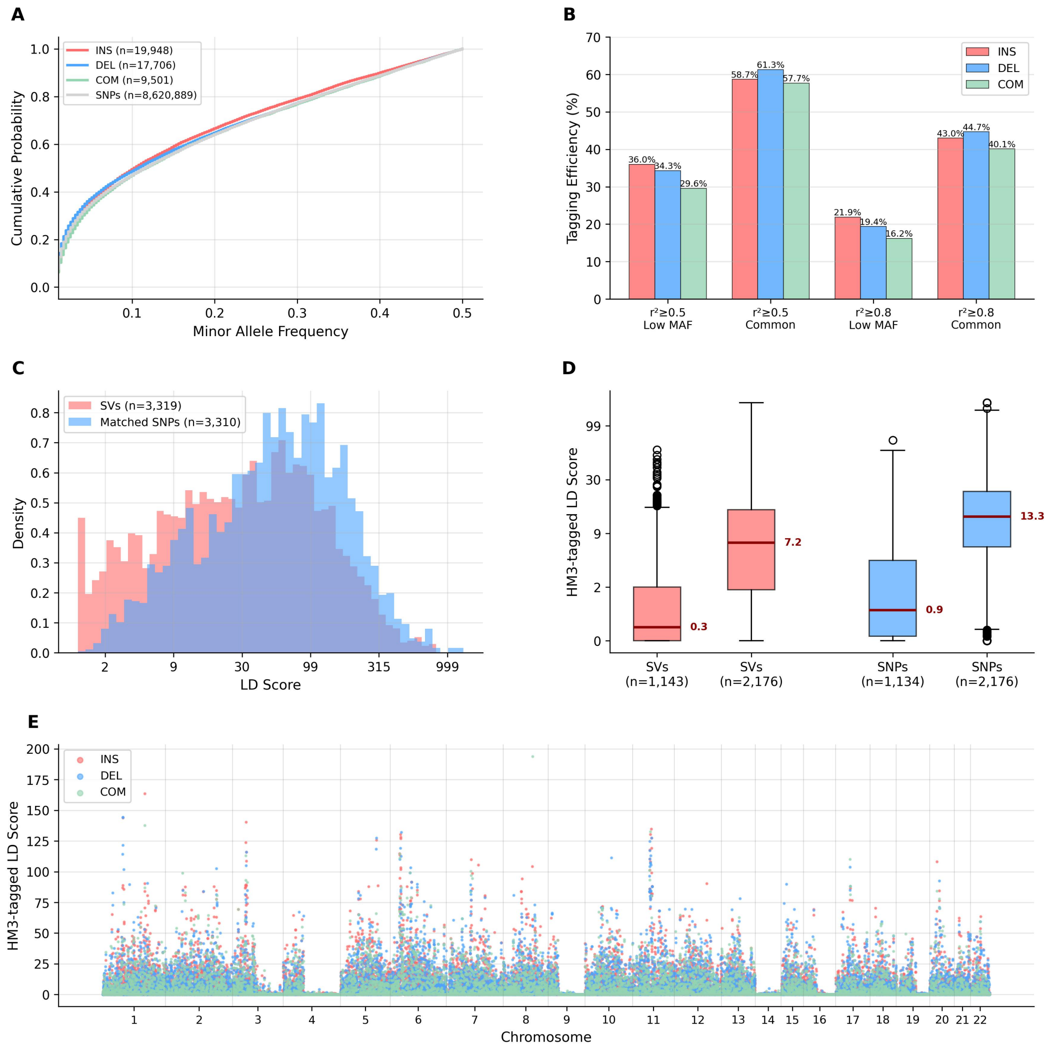


Supplementary Figure 1: **Structural variants exhibit substantial SNP tagging in East Asian ancestry. (A)** Cumulative MAF distributions show insertions enriched for rare alleles, while deletions and complex rearrangements resemble SNPs ($N$ = 47,155 SVs: 19,948 insertions, 17,706 deletions, 9,501 complex rearrangements). **(B)** SV tagging efficiency using all reference-panel SNPs as potential tags, stratified by MAF and $r^2$ threshold. Common SVs (MAF≥0.05) show 57.7 to 61.3% tagging at $r^2 \geq 0.5$, whereas low-MAF SVs show 29.6 to 36.0%. At the stringent threshold ($r^2 \geq 0.8$), common SVs show 40.1 to 44.7% tagging and low-MAF SVs 16.2 to 21.9%. **(C)** Reference LD score density distributions on chromosome 1 (SVs $n$ = 3,319, MAF-matched SNPs $n$ = 3,310) using all SNPs as tags, showing that SVs have somewhat lower but overlapping LD scores compared with SNPs. **(D)** HM3-tagged LD scores on chromosome 1 after restricting tag variants to HapMap3 SNPs, stratified by MAF (SVs low MAF $n$ = 1,143, SVs common $n$ = 2,176, SNPs low MAF $n$ = 1,134, SNPs common $n$ = 2,176). **(E)** Genome-wide HM3-tagged LD scores for all 47,155 SVs reveal LD hotspots and inter-chromosomal heterogeneity in SV tagging by HapMap3 SNPs.

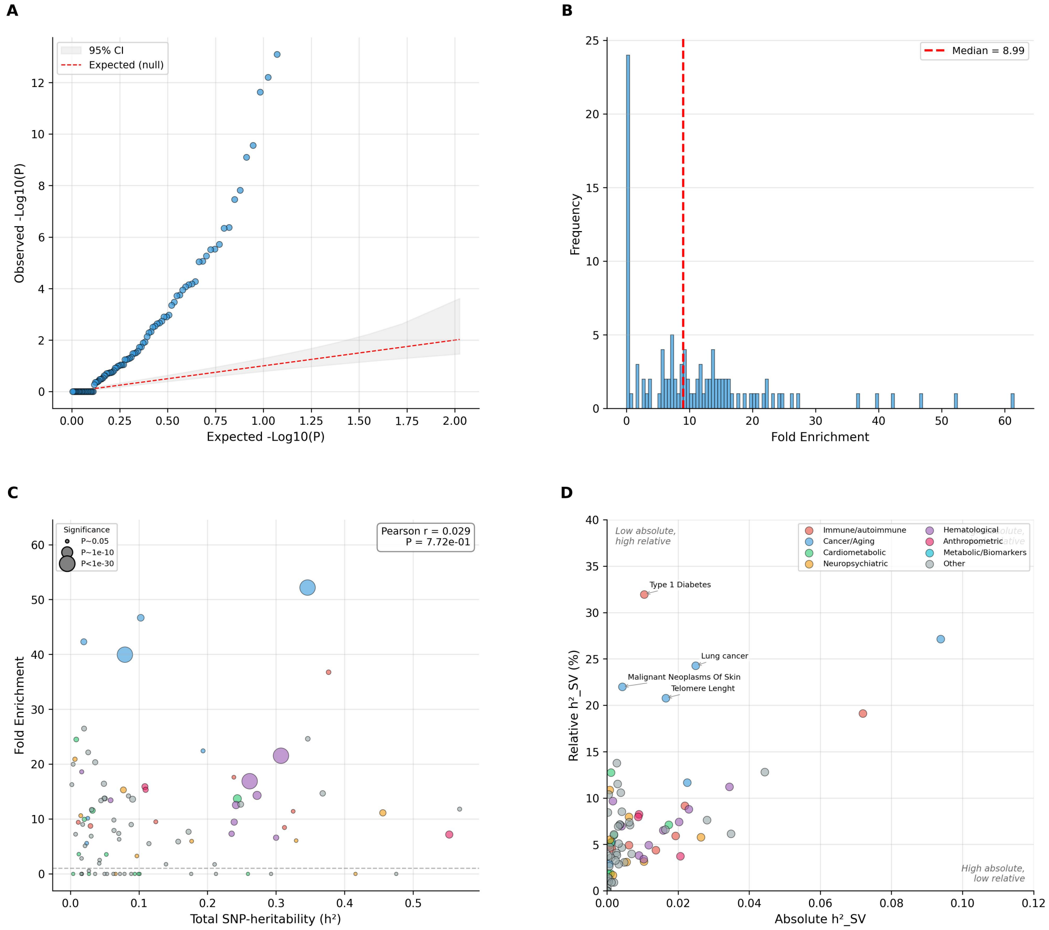


Supplementary Figure 2: **Extended analysis of SV enrichment patterns and heritability partitioning.** **(A)** Quantile-quantile plot of enrichment P-values across all 105 traits, comparing observed versus expected distributions under the null hypothesis of no enrichment. Strong deviation from the identity line (red dashed) indicates systematic enrichment signal beyond chance expectation. Gray shading represents 95% confidence interval for the null distribution. **(B)** Distribution of fold enrichment values across all 105 analyzed traits, showing a median of 8.99-fold (red dashed line) with a long tail of extreme effects concentrated in specific trait categories. **(C)** Scatter plot of fold enrichment versus total variant heritability for all 105 traits. Point sizes scale with statistical significance ($-\log_{10} P$); colors indicate trait category. Pearson correlation ($r = 0.029$, $P = 7.7 \times 10^{-1}$) demonstrates that enrichment is independent of total variant heritability, reflecting trait-specific genetic architecture rather than statistical power. **(D)** Absolute versus relative SV heritability contributions reveal distinct biological scenarios. X-axis shows absolute $h^2_{\mathrm{SV}}$; y-axis shows the percentage of total heritability from SVs. Type 1 diabetes exemplifies the top-left quadrant (low absolute, high relative: 32.0%), while anthropometric traits occupy the bottom-right (high absolute, low relative), reflecting their highly polygenic backgrounds.

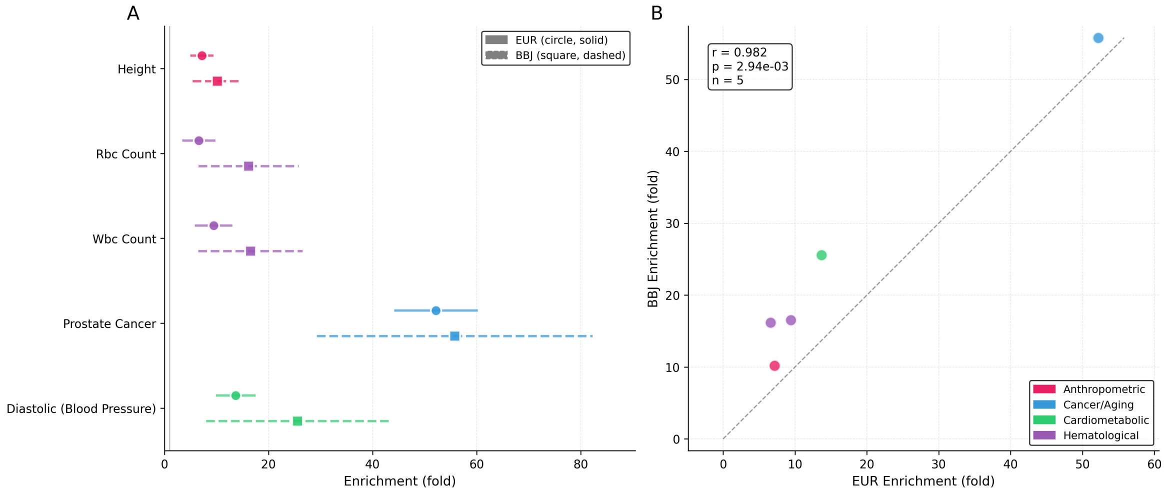


Supplementary Figure 3: **Cross-ancestry replication of SV enrichment in Biobank Japan. (A)** Overlapping forest plot comparing EUR discovery (circles, solid lines) and BBJ replication (squares, dashed lines) for five successfully replicated traits. Points are offset vertically for visibility. Error bars represent 95% confidence intervals. All five traits showed overlapping confidence intervals and statistically significant enrichment in both ancestries (Wald $P < 0.05$, LRT $P < 0.05$). **(B)** Correlation scatter plot demonstrating strong cross-ancestry consistency among replicated traits (Pearson $r = 0.982$, $P = 2.9 \times 10^{-3}$, $N = 5$). Dashed line represents perfect replication ($y = x$). Type 1 diabetes, lung cancer, and glaucoma failed replication criteria (see Supplementary Table S7 for complete results). Colors indicate trait categories.
